# Resplitting LIT-PCBA with MolSCISSOR: A Memory-Efficient, Quasi-Deterministic Framework for Reducing Structural and Physicochemical Biases

**DOI:** 10.64898/2026.11.26.769502

**Authors:** Cao-Minh Truong, Igor V. Tetko, Pedro J. Ballester, Olivier Taboureau, Viet-Khoa Tran-Nguyen

## Abstract

Reliable training-test partitioning is essential for developing and evaluating virtual screening (VS) methods, particularly in the era of machine learning (ML). However, existing molecular dataset splitting approaches do not simultaneously address key challenges in rigorous VS evaluation: cross-partition structural bias, test-set inter-class physicochemical bias, high memory requirements, limited scalability, stochastic allocation, a lack of automated optimization and of flexible training-test similarity control. We introduce `MolSCISSOR`, a lightweight, memory-efficient, quasi-deterministic framework for stratified data partitioning that integrates clustering and training-test allocation through multi-objective optimization. `MolSCISSOR` is used to repartition LIT-PCBA, a well-established VS benchmark comprising 15 target-specific datasets of experimentally confirmed actives and inactives, exhibiting substantial class imbalance, many of which are large-scale. We show that simple baseline ML models achieve significantly above-chance screening performance on the original `AVE`-derived partitions, indicating residual biases. The `MolSCISSOR`-generated benchmark, SPLIT-PCBA, is more challenging than both the `AVE` variant and that obtained using a `BitBIRCH`-Lean clustering-based workflow, with stricter cross-partition structural separation and improved physicochemical balance between test-set actives and inactives. `MolSCISSOR` mitigates stochastic variability, requires less memory than `AVE`, `Lo-Hi` and `DataSAIL`, and supports multiple split configurations with varying cross-partition similarity. `MolSCISSOR` and SPLIT-PCBA are freely available at https://github.com/caominhtr/molscissor and https://doi.org/10.5281/zenodo.23942580.

## 1 Introduction

Virtual screening (VS) of small molecules is a core task in computer-aided drug discovery. Screening power is commonly used to assess its performance, reflecting the ability of a VS procedure to prioritize and enrich active compounds among screened molecules [1]. Numerous benchmarking datasets have been introduced since the early 2000s for screening power assessment [2, 3, 4, 5, 6]. With the rapid development of artificial intelligence (AI), VS has increasingly moved beyond conventional approaches toward machine learning (ML)-based methods, raising the question of whether established benchmarks remain suitable for rigorously evaluating their predictive performance. Several studies have highlighted important structural and physicochemical biases in widely used legacy benchmarks such as DUD, DUD-E, and DEKOIS, which can artificially enrich true actives and inflate estimates of screening performance, notably of AI-based VS methods [7, 8, 9, 10, 11]. In response to these limitations, LIT-PCBA was introduced in 2020 as a more realistic and challenging benchmark for VS [12]. Unlike most predecessor datasets [7, 8, 9, 10, 11], LIT-PCBA consists entirely of experimentally confirmed active and inactive compounds with overlapping physicochemical spaces across 15 biologically diverse protein targets [12]. Its target-specific datasets also span a broad range of sizes, from relatively small (1,000-10,000 compounds) and medium-sized (10,000-100,000 compounds) to large (*>*100,000 compounds) collections. It further mimics the extreme class imbalance of real-world prospective screening campaigns, with active-to-inactive ratios as low as approximately 1:18,000 [12]. Consequently, LIT-PCBA has been widely adopted as an unbiased, realistic benchmark for modern ML-based scoring functions (SFs) and VS workflows [13, 14, 15, 16, 17, 18, 19, 20]. However, the full design of LIT-PCBA has not always been exploited. Indeed, most studies used the complete LIT-PCBA collection as a single evaluation dataset, some even further subset it according to their own *a priori* criteria to reduce the number of compounds and/or targets used for evaluation [10, 13, 15, 21]. Such non-standardized (and, in some cases, arbitrary) filtering retains only a portion of the benchmark, substantially alters its original composition, including class proportions, physicochemical and structural distributions, thereby undermining its design. Moreover, this approach is intended to evaluate the screening performance of generic VS workflows or SFs trained on external datasets such as DUD-E [4, 22] or PDBbind [23, 24, 25]. In these cases, retaining in LIT-PCBA exact or homologous targets and/or compounds related to those present in the external training data may introduce training-test contamination and artificially inflate screening power [26]. In fact, the field of AI-based VS in drug design has consistently highlighted the superior screening power of target-specific ML SFs over generic ones in both retrospective and prospective settings [27, 28, 29, 30, 31, 32]. This motivated the original LIT-PCBA design to provide, for each of its 15 target-ligand sets, two separate partitions specifically intended for training and testing target-specific ML SFs. Yet, this feature has remained largely underexploited, and it is unclear whether the original training-test partitions, generated using the Asymmetric Validation Embedding (`AVE`) method [11], provide optimal separation for this purpose or whether alternative splitting strategies could further improve upon `AVE`-derived partitions.

Introduced in 2018, `AVE` is a nearest-neighbor-based, similarity-aware approach that uses pairwise distances between compounds across activity classes in training and validation/test sets to quantify and minimize dataset bias based on structural fingerprints or molecular descriptors, simultaneously accounting for intra- and inter-class similarity across partitions [11]. However, AVE can become computationally intensive for large chemical libraries because of its pairwise distance calculations, motivating modifications to accelerate computation [12, 33]. Other recently introduced splitting approaches, such as Lo-Hi [34] and DataSAIL [35], have also been reported to face substantial memory demands when applied to large libraries, thus limiting their scalability in the era of large-scale datasets. Developments such as `BitBIRCH` for small-molecule clustering and its more recent implementation, `BitBIRCH-Lean (bblean)`, address this limitation by combining a Balanced Iterative Reducing and Clustering using Hierarchies (BIRCH)-like tree structure with the instant similarity formalism to process binary fingerprints more efficiently, substantially reducing computational costs and memory requirements during the molecular clustering stage of dataset splitting [36, 37]. However, both the clustering process itself and the eventual assignment of molecules to training and validation/test sets when implementing these clustering approaches may still be subject to stochasticity and residual biases, potentially compromising benchmark reproducibility and model generalization.

In this work, we demonstrate that many structural fingerprint-based, graph-based, and pre-trained transformer-based baseline models consistently achieve ROC-AUC (area under the receiver operating characteristic curve) values significantly above 0.5 on the original AVE-partitioned LIT-PCBA datasets. Thus, although the complete LIT-PCBA collection has been shown to pose a substantial challenge for screening power assessment, its AVE-derived training-test splits may still permit simple ML models to achieve screening performance significantly better than random selection. This indicates that residual biases may persist in partitions designed to be AVE-unbiased, underscoring the need for stricter, multi-objective partitioning frameworks to better assess model generalizability when LIT-PCBA partitions are used to train and evaluate target-specific ML SFs.

To this end, our study has four main objectives:

1. To introduce MolSCISSOR (**<u>Mol</u>**ecular **<u>S</u>**imilarity-aware **<u>C</u>**hemical **<u>I</u>**nter-class **<u>S</u>**tratified **<u>S</u>**plit for **<u>O</u>**bjective **<u>R</u>**epartitioning), a novel, memory-efficient, multi-objective optimization-based partitioning tool integrating molecular clustering, similarity constraints and physicochemical property matching with stratification to mitigate sampling bias in VS;
2. To construct and introduce SPLIT-PCBA, a new set of training-test partitions for LIT-PCBA generated using MolSCISSOR, designed to minimize structural and physicochemical biases, presenting a more rigorous benchmark for evaluating target-specific models in VS;
3. To systematically evaluate and benchmark three different data-splitting approaches on LIT-PCBA by quantifying structural bias alongside test-set artificial enrichment and predictive performance across benchmark variants, comparing `MolSCISSOR` directly with `AVE` [11] and a `MolSCISSOR`-equivalent splitting pipeline that uses bblean [36, 37], a modern, ultra-fast, high-throughput clustering algorithm designed for large chemical libraries;
4. To critically assess `AVE` and the `bblean` clustering-based splitting approach, particularly their susceptibility to stochastic variability and biases across training-test splits; and to demonstrate the memory inefficiency of two other recent splitting frameworks, `Lo-Hi` and `DataSAIL`, when applied to large-scale datasets comprising more than 100,000 molecules of LIT-PCBA.

The LIT-PCBA variants partitioned by `AVE`, the bblean clustering-based approach, and MolSCISSOR are hereafter referred to as `AVE`-unbiased LIT-PCBA, bblean-LIT-PCBA, and SPLIT-PCBA, respectively. All data and code are available on GitHub (https://github.com/caominhtr/molscissor) and Zenodo (https://doi.org/10.5281/zenodo.23942580).

## 2 Results

### 2.1 Working Principle of MolSCISSOR

The overall computational workflow of MolSCISSOR is illustrated in Figure 1. At its core, MolSCISSOR combines single-linkage clustering with a 0/1 knapsack-based allocation scheme. For each target, compounds are first grouped into disjoint clusters based on Tanimoto similarity between Morgan fingerprints (2,048 bits, radius 2) derived from RDKit-canonicalized SMILES strings, with active-and inactive-class compounds clustered separately. Entire clusters are subsequently assigned to output partitions via an iteratively updated allocation strategy. Actives first assigned greedily to the test partition serve as the initial reference set (*D*_ref_). Subsequently, inactive compounds are then allocated to match the DeepCoy property distributions of *D*_ref_, after which the newly allocated inactive set updates *D*_ref_ to guide a reciprocal re-allocation of active compounds. One complete round of allocating inactives relative to the reference active set, followed by a reciprocal re-allocation of actives, is defined as one cycle; by default, in our experiment, `MolSCISSOR` executes a single cycle to balance partitioning efficacy with computational efficiency. The partitioning decisions are governed by optimization using `Optuna` [38] (v.4.4.0) to tune the class-specific similarity used to define the clusters. In essence, MolSCISSOR executes multi-objective optimization to yield any training-to-test ratio between 3:1 and 4:1. The objectives simultaneously minimize both (i) maximum same-class Tanimoto similarity between training and test sets to prevent structural data leakage and bias, and (ii) test-set deviation from optimal embedding (DOE) scores to alleviate inter-class physicochemical separation or artificial enrichment. Non-dominated solutions along the Pareto front are evaluated, defining the final optimal split jointly by the selected pair of class-specific thresholds and the corresponding optimal training-test assignment obtained under those thresholds, with the trial yielding the lowest test-set DOE score chosen as the final output (cf. Methods).

**Fig. 1.**
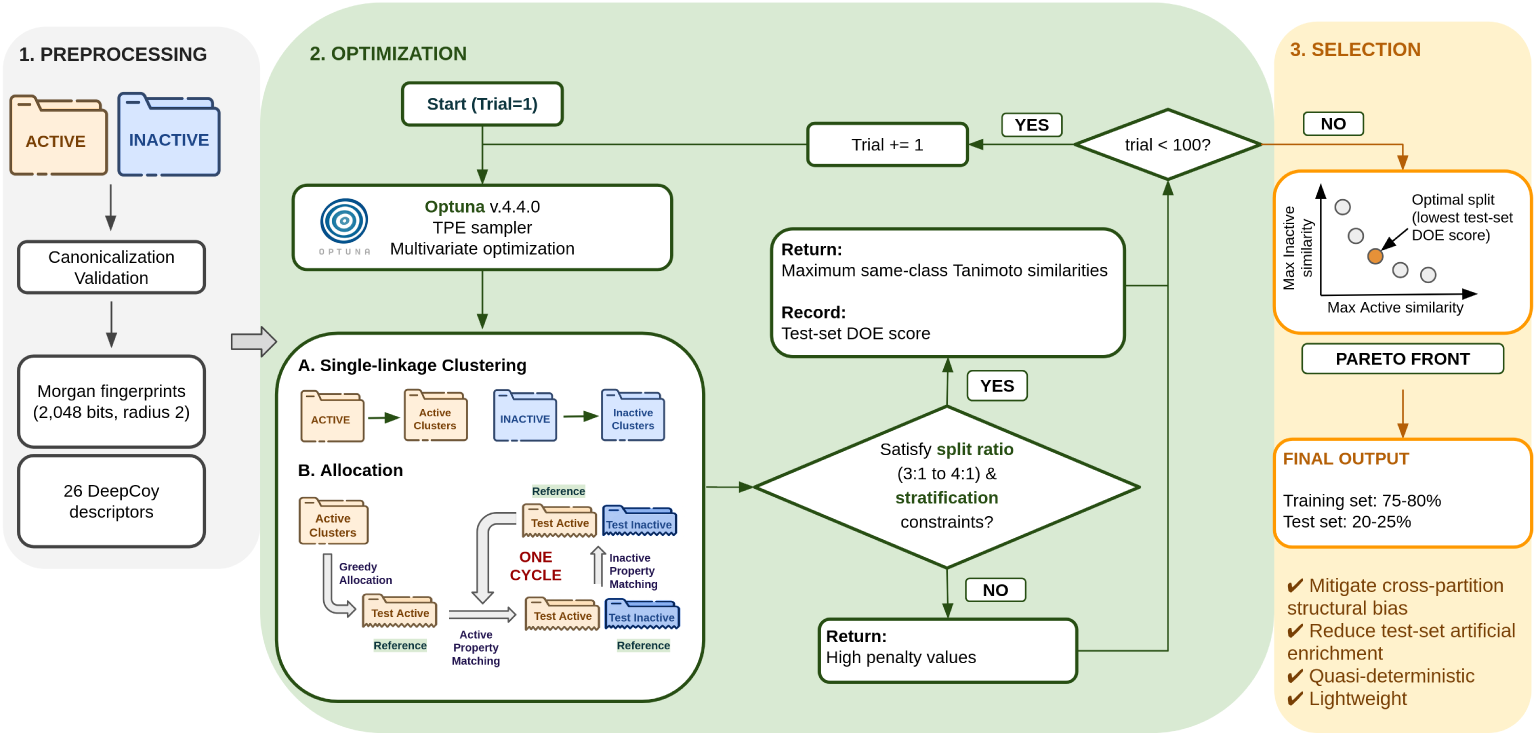
Overall computational workflow of `MolSCISSOR`. Step 1 (Preprocessing): SMILES strings of active and inactive compounds are canonicalized, validated, and encoded into Morgan fingerprints (2,048 bits, radius 2) alongside 26 DeepCoy physicochemical descriptors. Step 2 (Optimization): Optuna optimization is executed across 100 trials; in each trial, single-linkage clustering is applied independently to active and inactive compounds. An allocation scheme incorporating greedy allocation and iterative reciprocal re-allocation with updated reference sets yields a candidate training-test split, which is subsequently evaluated against target split-ratio and stratification constraints. Step 3 (Selection): Non-dominated solutions along the Pareto front are examined to select the single optimal split with the lowest test-set DOE score, producing the final partitioned dataset and the corresponding pair of class-specific similarity thresholds.

### 2.2 Simple machine-learning models achieve significantly above-chance screening performance on AVE-unbiased LIT-PCBA and bblean-LIT-PCBA, but not on MolSCISSOR-generated SPLIT-PCBA

The AVE-unbiased LIT-PCBA benchmark (https://drugdesign.unistra.fr/LIT-PCBA/, accessed March 2026) was initially designed to reduce dataset bias using the AVE splitting method, providing challenging training-test splits for target-specific ML-based models [11, 12]. Here, we trained and evaluated a series of baseline ML classifiers using default hyperparameters, including Random Forest (RF), Extreme Gradient Boosting (XGB), Support Vector Classifier (SVC), and Multi-Layer Perceptron (MLP) architecture from `DeepPurpose` [39] trained on Morgan fingerprints (2,048 bits, radius 2), Daylight fingerprints (2,048 bits) or Torsion fingerprints (2,048 bits). Additionally, we evaluated a Graph Convolutional Network (GraphConv) [40] operating on molecular graph representations, alongside a fine-tuned ChemBERTa transformer foundation model [41] pre-trained on 77 million compounds from PubChem. Each model configuration was trained across three random seeds. Three LIT-PCBA training-test variants were considered: AVE-unbiased LIT-PCBA, MolSCISSOR-derived SPLIT-PCBA, and partitions constructed from molecular clusters generated by bblean (bblean-LIT-PCBA). Since bblean is a clustering method rather than a complete training-test splitting tool, its clusters were assigned to the training or test set using the MolSCISSOR allocation strategy to ensure a consistent comparison between the two approaches, particularly their underlying clustering algorithms (cf. Methods). Across the 15 target partitions of the AVE-unbiased splits, the baseline models achieved mean ROC-AUCs from 0.581 to 0.709 on the test sets, significantly exceeding random performance (ROC-AUC *>* 0.5, one-sample two-sided Wilcoxon signed-rank test with Bonferroni correction, p-values *<* 0.001). An identical trend was observed on the bblean-LIT-PCBA variant, with mean ROC-AUCs from 0.568 to 0.650, significantly higher than 0.5, p-values *<* 0.01. These observations indicate that both splitting strategies retain biases sufficiently pronounced to be exploited by simple baseline models.

In contrast, the SPLIT-PCBA partitions produced by `MolSCISSOR` were substantially more challenging for all baseline models. Across the 15 target sets, mean test-set ROC-AUCs dropped to near the theoretical random baseline of 0.5, ranging from 0.490 to 0.553, and were statistically equivalent to random performance (one-sample two-sided Wilcoxon signed-rank test with Bonferroni correction, p-values *>* 0.05). This indicates that active compounds in the SPLIT-PCBA test partitions cannot be trivially distinguished from or prioritized over inactives by simple baseline models based on 2D molecular fingerprints or graph-based representations, or by a chemical foundation model. Interestingly, the relative ranking of the evaluated models remained consistent across the LIT-PCBA variants (Spearman’s rank correlation coefficient = 0.758 *−* 0.789, p-values *<* 0.01, Supplementary Data S1), supporting the robustness of retrospective model selection, an important consideration when prioritizing models for prospective VS campaigns [32]. Baseline model performances are summarized in Table 1, with detailed target-level data provided on GitHub and Zenodo (cf. Data availability).

**Table 1.** Predictive performance of baseline ML models on three LIT-PCBA training-test variants. Performance is reported as mean ± standard deviation of ROC-AUC values across 15 targets. Statistical comparisons were performed using the one-sample two-sided Wilcoxon signed-rank test against a threshold of 0.5 with Bonferroni correction. Only statistically significant comparisons are displayed with significance levels indicated as follows: * (0.01 < p-value ≤ 0.05), ** (0.001 < p-value ≤ 0.01), *** (0.0001 < p-value ≤ 0.001), and **** (p-value ≤ 0.0001).

| **Models** | **AVE-unbiased**<br>**LIT-PCBA** | **bblean-**<br>**LIT-PCBA** | **MolSCISSOR-generated**<br>**SPLIT-PCBA** |
|---|---|---|---|
| **Models employing Morgan fingerprints (2,048 bits, radius 2)** |  |  |  |
| RF | 0.583 ± 0.081<sup>****</sup> | 0.597 ± 0.094<sup>****</sup> | 0.508 ± 0.074 |
| XGB | 0.666 ± 0.098<sup>****</sup> | 0.645 ± 0.141<sup>****</sup> | 0.547 ± 0.144 |
| SVC | 0.681 ± 0.112<sup>****</sup> | 0.650 ± 0.142<sup>****</sup> | 0.544 ± 0.127 |
| MLP | 0.581 ± 0.104<sup>****</sup> | 0.604 ± 0.104<sup>****</sup> | 0.507 ± 0.120 |
| **Models employing Daylight fingerprints (2,048 bits)** |  |  |  |
| RF | 0.584 ± 0.086<sup>****</sup> | 0.568 ± 0.105<sup>***</sup> | 0.490 ± 0.084 |
| XGB | 0.651 ± 0.107<sup>****</sup> | 0.591 ± 0.146<sup>**</sup> | 0.522 ± 0.087 |
| SVC | 0.649 ± 0.139<sup>****</sup> | 0.591 ± 0.139<sup>***</sup> | 0.521 ± 0.098 |
| MLP | 0.642 ± 0.123<sup>****</sup> | 0.598 ± 0.151<sup>***</sup> | 0.543 ± 0.112 |
| **Models employing Torsion fingerprints (2,048 bits)** |  |  |  |
| RF | 0.594 ± 0.076<sup>****</sup> | 0.596 ± 0.119<sup>****</sup> | 0.500 ± 0.083 |
| XGB | 0.669 ± 0.104<sup>****</sup> | 0.631 ± 0.145<sup>****</sup> | 0.535 ± 0.123 |
| SVC | 0.661 ± 0.114<sup>****</sup> | 0.635 ± 0.138<sup>****</sup> | 0.531 ± 0.127 |
| MLP | 0.587 ± 0.117<sup>***</sup> | 0.613 ± 0.109<sup>****</sup> | 0.517 ± 0.124 |
| **Graph-based model** |  |  |  |
| GraphConv | 0.633 ± 0.125<sup>****</sup> | 0.620 ± 0.114<sup>****</sup> | 0.544 ± 0.134 |
| **Fine-tuned foundation model** |  |  |  |
| ChemBERTa-77M-MTR | 0.709 ± 0.110<sup>****</sup> | 0.644 ± 0.176<sup>****</sup> | 0.553 ± 0.163 |

### 2.3 SPLIT-PCBA offers a more challenging virtual screening benchmark through improved bias control by MolSCISSOR

The difficulty of the screening task across different LIT-PCBA split variants was systematically assessed using the same baseline models (RF, XGB, SVC, MLP, GraphConv, and ChemBERTa) and three screening power metrics: ROC-AUC, NEF1% (normalized enrichment factor of true hits at the top 1%-ranked molecules), and BEDROC (Boltzmann-enhanced discrimination of receiver operating characteristic) score with *α* = 160.9 (corresponding to 80% of the score contribution arising from the top 1% of the ranked list [42]). Under this framework, a split variant is considered more challenging if models trained on its training partition achieve lower predictive performance on the corresponding test partition than those trained and evaluated on alternative splits, with all other settings held constant, including model hyperparameters, features, and evaluation protocols (cf. Methods). Results across the three LIT-PCBA variants are summarized in Figure 2, with raw data provided on GitHub and Zenodo (cf. Data availability).

**Fig. 2.**
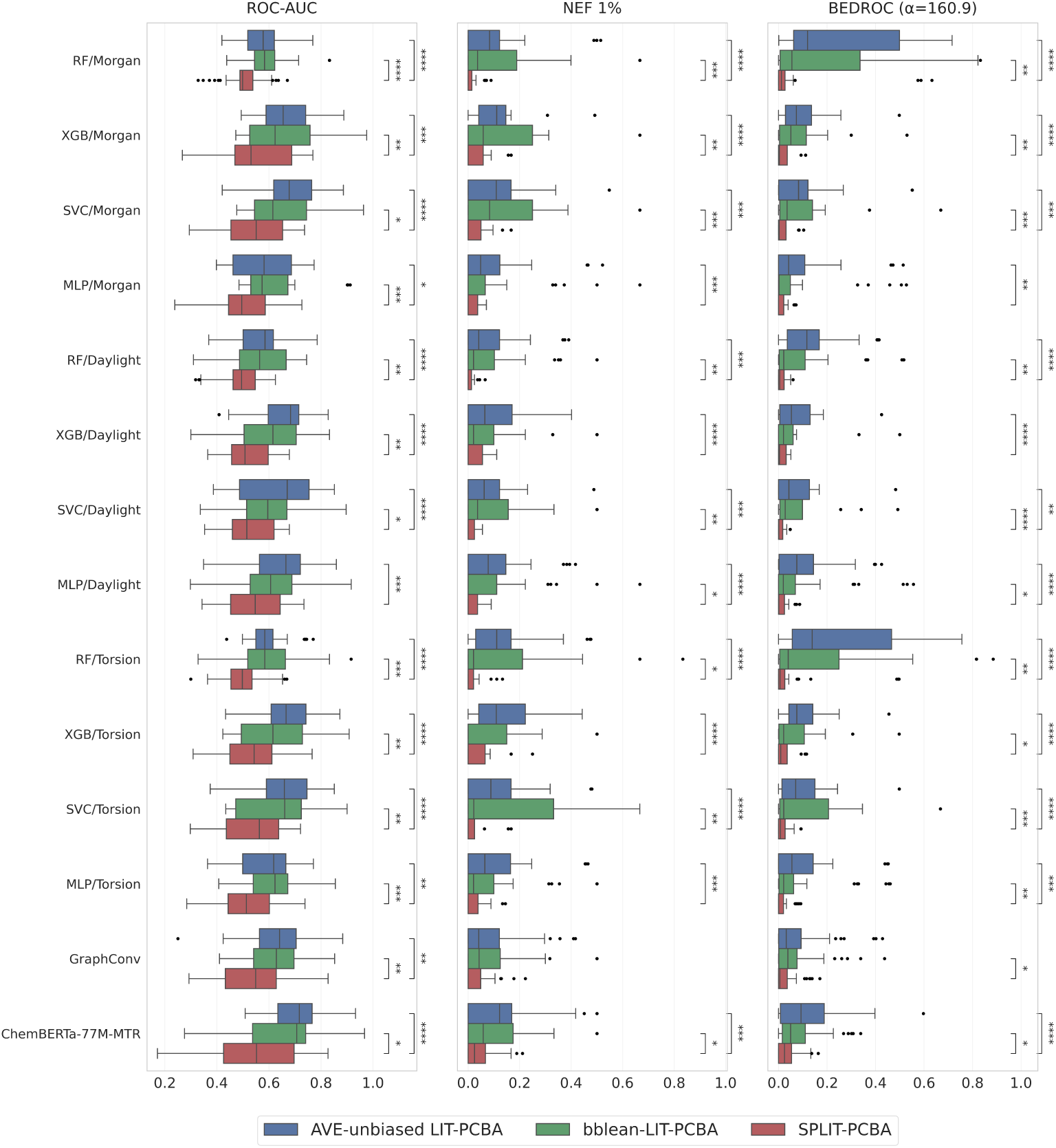
Comparison of baseline ML models (RF, XGB, SVC, and MLP) using molecular fingerprints, including Morgan fingerprints (2,048 bits, radius 2), Daylight fingerprints (2,048 bits), and Torsion fingerprints (2,048 bits), together with GraphConv and ChemBERTa-77M-MTR models across AVE-unbiased LIT-PCBA (blue), bblean-LIT-PCBA (green), and SPLIT-PCBA (red). Box plots show the distributions of screening performance across targets, with the horizontal line inside each box representing the median, and filled circles denoting outliers defined using the 1.5IQR rule, quantified by ROC-AUC (left panels), NEF1% (middle panels), and BEDROC(*α* = 160.9) (right panels). Statistical comparisons were performed using the two-sided Mann-Whitney U test with Bonferroni correction. Only statistically significant comparisons are displayed with significance levels indicated as follows: * (0.01 *<* p-value *≤* 0.05), ** (0.001 *<* p-value *≤* 0.01), *** (0.0001 *<* p-value *≤* 0.001), and **** (p-value *≤* 0.0001).

As shown in Figure 2, all baseline models, including fingerprint-based, graph-based, and pre-trained transformer-based models, achieved substantially lower test-set performance on SPLIT-PCBA than on the AVE-unbiased and bblean variants, highlighting the greater challenge posed by `MolSCISSOR`-derived partitions. By challenging both overall ranking performance (ROC-AUC) and early active-compound retrieval (NEF1% and BEDROC with *α* = 160.9, which are more important in practical VS campaigns [42, 43, 44]), SPLIT-PCBA offers a more stringent benchmark for assessing target-specific model generalizability. This is particularly relevant to real-world prospective screening campaigns involving medium-to large-scale compound libraries, where only a small number of true hits are typically identified [12, 45, 46, 47, 48].

#### 2.3.1 MolSCISSOR reduces cross-partition structural similarity, AVE bias, and scaled DataSAIL information “leakage”

To investigate why `MolSCISSOR`-generated SPLIT-PCBA partitions pose a greater challenge to baseline models than those derived from AVE and bblean, we first quantified the structural dissimilarity between the training and test sets across all split variants. Modern ultra-large chemical libraries encompass billions or even trillions of accessible compounds [49, 50], making chemical diversity increasingly important for broad target coverage and novel hit identification [51]. Accordingly, VS benchmarks should minimize structural similarity across training and test sets to assess model generalizability beyond closely related compounds, and better reflect realistic applications [11], particularly scaffold hopping [52, 53, 54]. In this context, we characterized structural bias using three key metrics: (i) intra-class cross-partition maximum similarity (Eq. (1)), which quantifies structural near-neighbor bias between same-class training- and test-set molecules; (ii) AVE bias (Eq. (2)) [11], which measures chemical space separation across the four subsets: training actives, training inactives, test actives, and test inactives; and (iii) scaled DataSAIL information “leakage” (scaled L(*π*), Eq. (3)) [35], according to DataSAIL’s authors, which evaluates the proportion of total dataset similarity that exists across partitions. The maximum Tanimoto similarities calculated using Morgan fingerprints (2,048 bits, radius 2) between training and test sets for the active (maxActive) and inactive (maxInactive) classes, alongside AVE bias and scaled L(*π*), are summarized in Figure 3, with raw data provided in Supplementary Table S1.

**Fig. 3.**
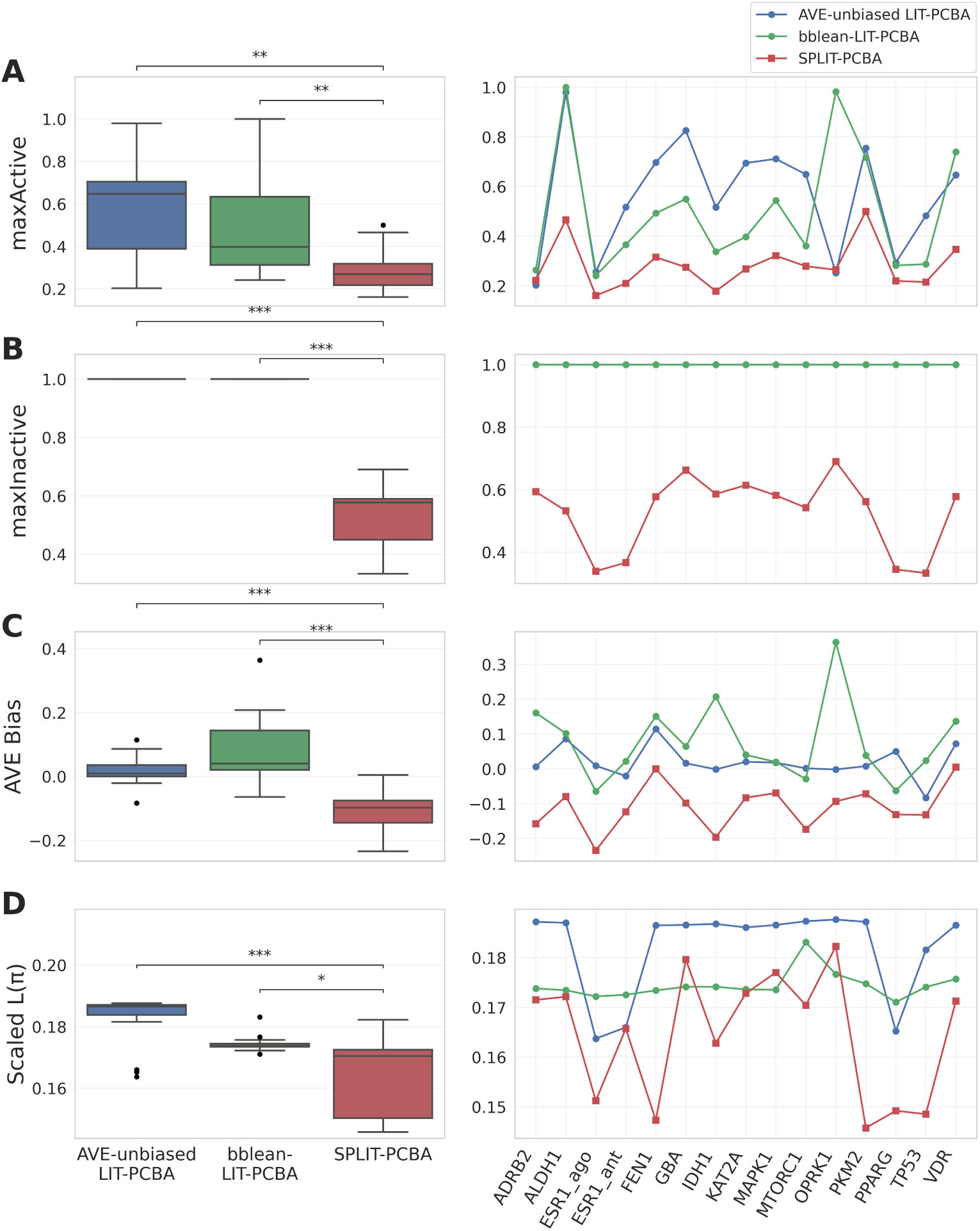
Comparison of maxActive (A), maxInactive (B), AVE bias (C), and scaled L(*π*) (D) across AVE-unbiased LIT-PCBA (blue), bblean-LIT-PCBA (green), and SPLIT-PCBA (red). Overall distributions across targets are presented as box plots (left panels) with the horizontal line inside each box representing the median, and filled circles denoting outliers defined using the 1.5IQR rule, alongside target-level values for 15 individual targets (right panels). Statistical comparisons were performed using the two-sided Mann-Whitney U test with Bonferroni correction. Only statistically significant comparisons are displayed with significance levels indicated as follows: * (0.01 *<* p-value *≤* 0.05), ** (0.001 *<* p-value *≤* 0.01), *** (0.0001 *<* p-value *≤* 0.001), and **** (p-value *≤* 0.0001).

In SPLIT-PCBA, cross-partition maximum Tanimoto similarities (Morgan fingerprints, 2,048 bits, radius 2) for both active and inactive classes were significantly lower than those observed in the other two benchmark variants (two-sided Mann-Whitney U test with Bonferroni correction, p-values *<* 0.01; Figure 3A, B). Notably, the maximum similarity for the inactive class was 1.000 across all 15 targets in both AVE-unbiased LIT-PCBA and bblean-LIT-PCBA, indicating that identical or near-identical same-class analogs, such as stereoisomers and congeneric compounds, were partitioned across training and test sets and, in the case of bblean, assigned to different clusters during the preceding clustering step. Similarly, both variants contained active compounds with near-perfect cross-partition similarity. For example, the ALDH1 target set in AVE-unbiased LIT-PCBA exhibited a maxActive = 0.979, while the OPRK1 target set in bblean-LIT-PCBA had a maxActive = 0.982. Such substantial 2D structural overlap introduces a bias that encourages model memorization, artificially inflating estimates of screening performance via near-neighbor retrieval. In contrast, by combining single-linkage clustering with optimization to maximize same-class structural dissimilarity between training and test sets under dual stratification constraints (cf. Methods), MolSCISSOR successfully reduces this bias, yielding markedly lower cross-partition maximum similarity for both classes (maxActive = 0.283 *±* 0.097 and maxInactive = 0.527 *±* 0.120). This greater structural separation challenges target-specific VS models to generalize to chemotypes that differ structurally from those encountered during training.

Additionally, SPLIT-PCBA exhibited a consistently lower AVE bias compared to the other two variants across all 15 targets (Figure 3C). This global reduction was statistically significant (two-sided Mann-Whitney U test with Bonferroni correction, p-values *<* 0.001), confirming that the bias mitigation was pronounced and consistent. Interestingly, MolSCISSOR does not directly minimize the AVE bias metric, unlike the genetic algorithm proposed by Wallach and Heifets [11]. Indeed, the standard AVE bias formulation calculates the net difference between same-class and inter-class structural similarity terms, a quantity not explicitly incorporated into the objective function of MolSCISSOR (cf. Methods). Furthermore, target-level analysis (Supplementary Table S1) revealed that only two out of 15 MolSCISSOR-repartitioned target-ligand sets exhibited slightly positive AVE bias scores (FEN1 at 0.0003 and VDR at 0.0049, both near zero), while the remaining targets yielded negative values. By comparison, nearly half of the 15 AVE-unbiased LIT-PCBA target sets did not reach the target AVE bias of below 0.01, illustrating that the AVE partitioning strategy does not guarantee convergence to the global optimum, as acknowledged by its authors [11]; while the training-test partitions obtained by allocating bblean clusters using the same allocation strategy as MolSCISSOR exhibited even higher AVE bias, exceeding 0.01 for 12 targets and even greater than 0.10 for some of them. The predominantly negative AVE bias values obtained with MolSCISSOR indicate that same-class separation across the training-test split exceeds inter-class separation, meaning training actives share greater structural similarity with test inactives than with test actives and vice versa, suggesting reduced cross-partition structural redundancy. An identical trend was observed for the scaled L(*π*) metric introduced by DataSAIL’s authors (Figure 3D), where SPLIT-PCBA achieved a significantly lower “leakage score” compared to the other two variants (two-sided Mann-Whitney U test with Bonferroni correction, p-values *<* 0.05). By globally minimizing intra-class cross-partition similarities across the entire dataset, MolSCISSOR suppresses total information leakage and alleviates sampling bias, further highlighting the robustness of our framework in preventing models from achieving inflated VS performance through structural memorization.

#### 2.3.2 MolSCISSOR better controls artificial hit enrichment driven by test-set inter-class physicochemical bias

In addition to maximizing the structural distance between same-class molecules across training and test sets, the multi-objective optimization implemented in MolSCISSOR is designed to reduce physicochemical discrepancies between actives and inactives within the test set. To assess this, we trained and evaluated additional RF, XGB, SVC, and MLP classifiers using 217 RDKit physicochemical descriptors as features on the corresponding training-test partitions of AVE-unbiased LIT-PCBA, bblean-LIT-PCBA, and SPLIT-PCBA. Screening power was evaluated using ROC-AUC, NEF1%, and BEDROC(*α* = 160.9). The results are summarized in Figure 4 and raw data are provided on GitHub and Zenodo (cf. Data availability).

**Fig. 4.**
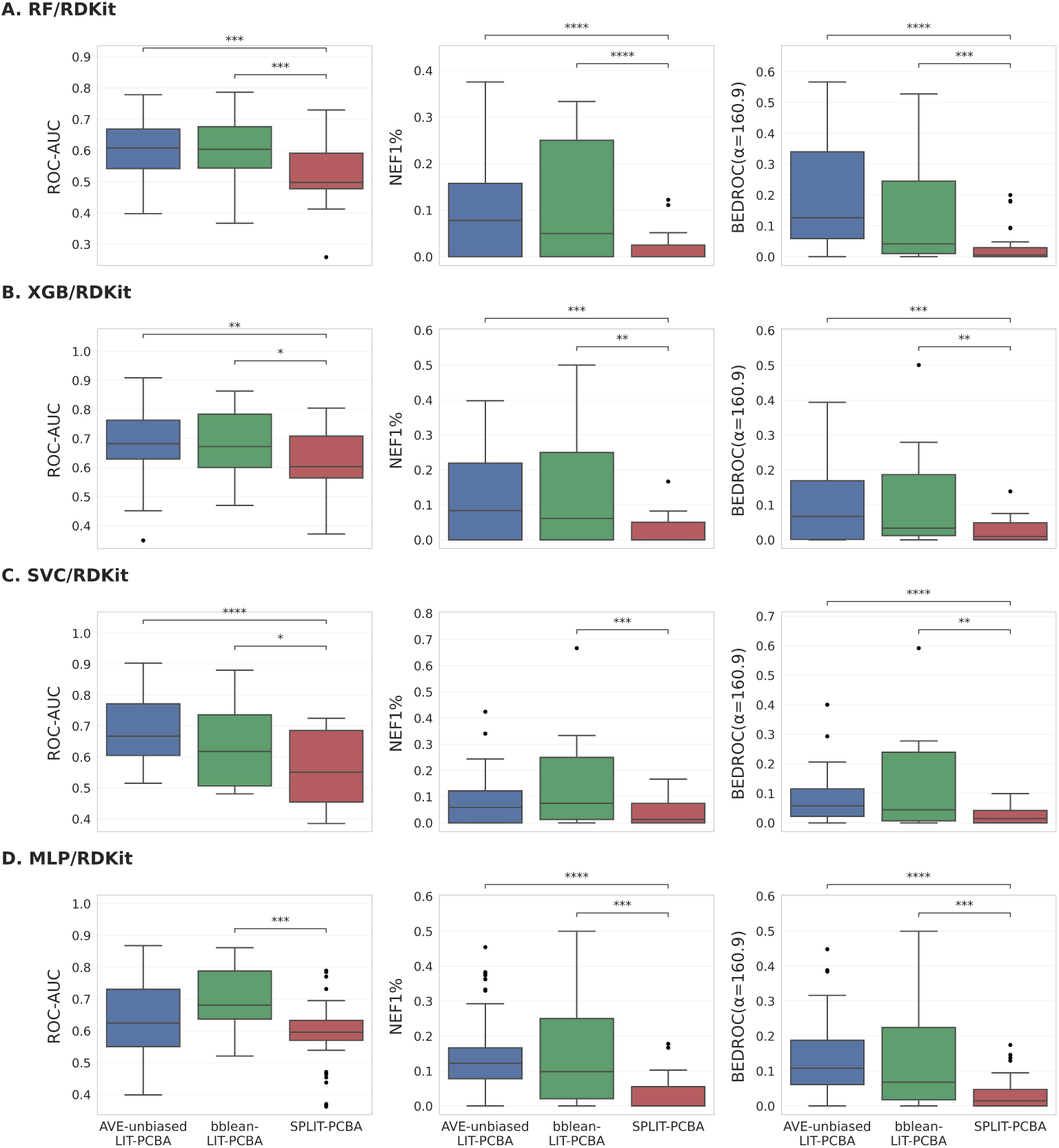
Comparison of additional ML models, RF (A), XGB (B), SVC (C), and MLP (D), each using 217 RDKit physicochemical descriptors, across AVE-unbiased LIT-PCBA (blue), bblean-LIT-PCBA (green), and SPLIT-PCBA (red). Box plots show the distributions of screening performance across targets, with the horizontal line inside each box representing the median, and filled circles denoting outliers defined using the 1.5IQR rule, quantified using ROC-AUC (left panels), NEF1% (middle panels), and BEDROC(*α* = 160.9) (right panels). Statistical comparisons were performed using the two-sided Mann-Whitney U test with Bonferroni correction. Only statistically significant comparisons are displayed with significance levels indicated as follows: * (0.01 *<* p-value *≤* 0.05), ** (0.001 *<* p-value *≤* 0.01), *** (0.0001 *<* p-value *≤* 0.001), and **** (p-value *≤* 0.0001).

By using 217 RDKit physicochemical descriptors as input features for these models instead of the 26 DeepCoy descriptors employed by MolSCISSOR during its allocation process, we ensured that the assessment was independent of the optimized variables, capturing a much broader physicochemical space. Even under this independent evaluation, models relying on physicochemical descriptors exhibited significantly poorer test-set screening power on SPLIT-PCBA than on the AVE-unbiased and bblean variants. This suggests that MolSCISSOR more effectively limits the ability of such models to exploit inter-class differences within the test set, thereby mitigating artificial enrichment of true actives and supporting the effectiveness of its allocation strategy.

### 2.4 MolSCISSOR demonstrates superior memory efficiency and scalability for medium- and large-sized datasets compared with AVE, Lo-Hi, and DataSAIL

Regarding practical utility, MolSCISSOR offers a lightweight and efficient data-splitting framework. While its memory efficiency is comparable to that of bblean, a state-of-the-art clustering tool designed for large chemical libraries, `MolSCISSOR` significantly outperforms the AVE, Lo-Hi, and DataSAIL methods in memory efficiency and scalability under identical execution conditions and using default parameter settings (cf. Methods). The AVE method imposes severe memory bottlenecks on large datasets by precomputing three dense pairwise distance matrices (active-active, inactive-inactive, and active-inactive). This yields a quadratic memory footprint *O*(*N* ^2^) that can easily consume hundreds of gigabytes (GB) of RAM when processing datasets exceeding 100,000 compounds. Although AVE supports multi-core parallelization via worker processes (-numWorkers), passing these massive distance matrices to worker threads introduces substantial communication overhead and reduces parallel efficiency. Similar computational and memory scaling challenges are also observed in other recently published advanced partitioning algorithms designed to mitigate information leakage and sampling bias, such as DataSAIL [35] and Lo-Hi [34] splitters. Specifically, since DataSAIL relies on precomputed pairwise similarity matrices coupled with scikit-learn’s agglomerative clustering, and Lo-Hi constructs explicit dense graph representations via NetworkX, both approaches become memory-intensive when applied to large screening libraries. A detailed comparison of these memory footprints (Supplementary Table S2) underscores MolSCISSOR’s ability to bypass these bottlenecks. Indeed, for medium-sized datasets (10,000-100,000 compounds per set), MolSCISSOR requires only 1.0-1.5 GB of RAM, comparable to bblean, whereas AVE, Lo-Hi, and DataSAIL require 10-200 GB. The difference becomes even more pronounced for large datasets containing more than 100,000 compounds each. `MolSCISSOR` and bblean require no more than 8.0 GB of memory, even when splitting the largest LIT-PCBA dataset (IDH1, nearly 360,000 molecules). In contrast, AVE, Lo-Hi, and DataSAIL frequently fail to complete the splitting process under a 200 GB RAM limit because of Out-Of-Memory errors, with some jobs, notably half of those involving DataSAIL, remaining infeasible even with 1 TB of RAM. These observations establish MolSCISSOR as a scalable and memory-efficient solution for dataset partitioning, particularly suited to large datasets.

## 3 Discussion

This study presents SPLIT-PCBA, a repartitioned variant of the LIT-PCBA dataset generated by MolSCISSOR, offering a more rigorous alternative to the original AVE-unbiased partitions and another variant constructed by allocating molecular clusters generated by bblean (bblean-LIT-PCBA). On SPLIT-PCBA, target-specific fingerprint-based, graph-based, and pre-trained transformer-based models achieved lower performance in both global ranking discrimination and early hit retrieval. This systematic decline in screening power is consistent with the more effective reduction of both cross-partition structural bias and inter-class physicochemical bias within the test set achieved by MolSCISSOR. Such well-documented biases in VS reward the trivial memorization of near-neighbor chemotypes and facilitate the discrimination of test-set ligands belonging to different activity classes [11, 21, 25, 27, 55], as observed with scaffold-based splitting, an approach widely adopted in community benchmarks such as MoleculeNet [56, 57]. MolSCISSOR addresses these biases by enforcing class-specific similarity thresholds and optimizing post-clustering ligand allocation. This optimization promotes structural separation between same-class compounds across partitions while making inter-class similarity outweigh intra-class similarity, as reflected by significantly lower maxActive, maxInactive, `AVE` bias, and scaled L(*π*) values. Notably, neither AVE nor bblean explicitly accounts for physicochemical bias: their resulting test partitions may therefore retain inter-class differences in these properties that ML models can exploit to distinguish actives from inactives more easily. By contrast, MolSCISSOR uses the test-set DOE, which addresses this source of bias [2, 3, 58], as an explicit tie-breaking criterion for selecting the final partition from the Pareto front generated by the training-test allocation process [59]. Its iterative refinement further corrects residual spatial mismatches by using one class as a reference profile when adjusting the allocation of the other, thereby promoting physicochemical balance between test-set actives and inactives. Together, molecular clustering and training-test allocation enable `MolSCISSOR` to address VS biases within a unified workflow.

A critical consideration when selecting a data partitioning framework is its intrinsic algorithmic stochasticity. MolSCISSOR adopts a quasi-deterministic approach that more rigorously controls stochastic variability than alternative clustering-based splitting methods, owing to two primary design choices. First, its single-linkage clustering generates strictly deterministic disjoint clusters for any given similarity threshold by identifying connected components based on graph connectivity, without random initialization or centroid selection [60, 61]. Unlike k-means [62] or Butina [63], it avoids convergence to different local optima and seed-dependent variability. In contrast, the recently published UMAP-based approach combined with Ward’s linkage [44] introduces stochasticity through UMAP’s probabilistic embedding, seed selection, hyperparameter choices and tuning, while also requiring the number of clusters to be predefined. Although bblean avoids random seeds, its hierarchical Clustering Feature tree introduces substantial data-order dependency: sequential assignment of molecules to dynamically updated cluster centroids makes the resulting cluster boundaries sensitive to input ordering [36, 37]. This dependence, together with user-defined hyperparameters such as branching factor and merging criterion, compromises reproducibility, as further illustrated in Supplementary Data S2. Second, MolSCISSOR constrains stochasticity during allocation through three mechanisms: (i) dual stratification constraints maintaining approximately the same active-to-inactive ratio in both subsets and enforcing the user-defined training-to-test ratio range; (ii) a greedy allocation strategy followed by deterministic 0/1 knapsack-based optimization [59]; and (iii) iterative refinement based on predefined update rules rather than random reassignment. By contrast, AVE relies on random initialization of its candidate population and probabilistic mutation, through which compounds are arbitrarily added or dropped. Neither its original implementation [11] nor the Python 3-compatible version provided in a previous study [43] sets global random seeds. Consequently, repeated executions on identical inputs can yield different split compositions and subset sizes, complicating cross-study reproducibility.

Another distinguishing feature of `MolSCISSOR` is its memory efficiency and scalability compared with recent dataset partitioning approaches such as `Lo-Hi` and `DataSAIL`. In their original studies, these methods were evaluated on relatively small- to medium-sized molecular datasets drawn from MoleculeNet, Therapeutics Data Commons and MolData, most containing fewer than 10,000 compounds and none exceeding 50,000 [34, 35]. Such datasets do not fully reflect the scale of contemporary chemical collections, which increasingly include large- and ultra-large-scale datasets [12, 29, 45]. Computational feasibility is therefore an important consideration for rigorous model training and evaluation. LIT-PCBA provides a more demanding setting for assessing partitioning scalability, with over half of its VS datasets containing more than 100,000 compounds. In this setting, MolSCISSOR requires considerably less memory than Lo-Hi and DataSAIL for medium-sized datasets and can partition large datasets that these methods often fail to process even with 1 TB of RAM. Its lower memory footprint consequently reduces computational requirements and facilitates partitioning large chemical collections in resource-constrained environments. Another practical advantage of MolSCISSOR is its automated selection of class-specific molecular similarity thresholds. Unlike Lo-Hi and DataSAIL, which rely on a predefined threshold or number of clusters to group actives and inactives together, MolSCISSOR uses Optuna-based optimization to identify suitable thresholds independently for each activity class. The fixed settings in the former two tools may produce clusters too large for the desired training-test partition sizes, preventing successful allocation and potentially requiring manual modification of thresholds or training-to-test ratios in the source code. MolSCISSOR instead identifies the lowest feasible class-specific similarity thresholds that permit allocation within the user-defined ratio range. This data-dependent optimization avoids imposing a single set of thresholds across datasets with different chemical compositions and class distributions (as in Lo-Hi and DataSAIL), or a predefined formula for determining such thresholds (as in bblean), while promoting greater cross-partition structural separation. This added optimization capability comes, as expected, with additional computational time, which only becomes noticeable for large-scale datasets: splitting those containing over 100,000 molecules can require more than five days, whereas small- and medium-sized datasets generally require only minutes to a few hours under our default execution conditions (cf. Benchmarking memory footprints). MolSCISSOR therefore offers an efficient, automated tool for VS benchmarking, without requiring users to modify the source code or manually adjust clustering parameters through trial and error.

In addition to its quasi-deterministic operation and automated threshold optimization, MolSCISSOR also allows users to define their own class-specific similarity thresholds. By default, MolSCISSOR activates the Optuna-based tuning described above and generates a corresponding training-test split for a given dataset. Alternatively, users can disable this optimization using the -no tune flag and specify their preferred thresholds through the -thresholdActive and -thresholdInactive arguments. Different threshold pairs yield different cluster configurations, which are then allocated to the training and test sets using the same allocation strategy. This flexibility permits users to generate multiple splits with varying levels of same-class, cross-partition structural similarity and systematically assess model generalizability under different training-test similarity conditions (i.e., across multiple splits spanning different levels of training-test dissimilarity). Thus, beyond supporting more rigorous assessment through reduced structural and physicochemical biases (which are recognized challenges in VS benchmarking), MolSCISSOR also enables similarity-spectrum-based evaluation [64]. Performance metrics and degradation curves can then characterize how predictive performance changes as the test compounds become less similar to those in the training set, providing insights into the extent to which patterns learned from the training data remain predictive for less similar compounds [64, 65, 66]. Moreover, MolSCISSOR can integrate user-provided clustering results into its allocation workflow, in addition to its built-in single-linkage clustering. As demonstrated in this study, the bblean clustering approach has been integrated, illustrating the workflow’s compatibility with alternative clustering methods. Users can further employ cluster assignments generated using any clustering algorithm of their choice by providing input datasets containing the corresponding cluster indices and specifying the `-algo other` flag instead of -algo molscissor or -algo bblean (see our GitHub repository; cf. Code availability). Taken together, MolSCISSOR combines memory efficiency, lightweight optimization of class-specific similarity thresholds, quasi-deterministic allocation of molecular clusters, mitigation of biases that can inflate screening performance, user-defined control over training-test similarity, and compatibility with externally generated clustering results. It supports both the generation of a single split for assessing model generalizability in VS and systematic evaluation across a spectrum of similarity levels, helping users interpret model performance in relation to the structural separation represented by each split.

## 4 Methods

### 4.1 Data retrieval and preprocessing

The AVE-unbiased [11] LIT-PCBA dataset was retrieved from the official repository (https://drugdesign.unistra.fr/LIT-PCBA/, accessed March 2026), including four predefined subsets for each target: training actives, training inactives, test actives, and test inactives (here, “validation” in AVE terminology refers to the test set). We recombined these four subsets into two: all actives and all inactives. These combined sets were then fed directly into our MolSCISSOR workflow or processed using a MolSCISSOR-equivalent partitioning procedure that employed bblean for clustering, resulting in the SPLIT-PCBA and bblean-LIT-PCBA variants, respectively. Compounds were represented as SMILES strings, which were then canonicalized and validated via 2D structure generation using AddMoleculeColumnToFrame from rdkit.Chem.PandasTools (RDKit, v.2025.3.3). For MolSCISSOR, the resulting RDKit molecule objects were directly used to generate Morgan fingerprints (2,048 bits, radius 2) using GetMorganFingerprintAsBitVect from rdkit.Chem.AllChem. In addition, these molecule objects were used to calculate 26 DeepCoy physicochemical descriptors [58] using the decoy utils.py script provided in the official DeepCoy GitHub repository (https://github.com/fimrie/DeepCoy/blob/master/evaluation/decoy_utils.py, accessed March 2026). The complete list of these descriptors is detailed in Supplementary List S1. These descriptors were then standardized using scikit-learn’s StandardScaler (v.1.7.2). The Morgan fingerprints were generated to compute Tanimoto similarities as thresholds for linkage clustering, whereas the DeepCoy descriptors were utilized to calculate DOE scores, serving as a tie-breaking criterion to select the best solutions from the Pareto front during the optimization process (cf. Core algorithm). For bblean-based split, the canonicalized and validated SMILES strings were directly used for clustering, while the allocation of disjoint clusters into training and test partitions followed the same protocol used in MolSCISSOR based on Euclidean distances between 26-dimensional DeepCoy physicochemical descriptor vectors. This ensures a framework equivalent to MolSCISSOR in terms of partition allocation and differing only in the clustering method (cf. Allocation strategy & cf. bblean clustering-based split).

### 4.2 Core algorithm

MolSCISSOR is presented herein as an optimization-based dataset partitioning tool designed to optimize training-test splitting, utilizing Optuna (v.4.4.0) [38] as the core optimization engine. Specifically, MolSCISSOR performs multivariate optimization to simultaneously minimize the maximum pairwise Tanimoto similarity using Morgan fingerprints (2,048 bits, radius 2) between compounds of the same class across the training and test sets (Algorithm 1). In other words, our tool solves for the minimal threshold of maximum pairwise Tanimoto similarity for each class required to partition a dataset while satisfying the dual constraints of stratified sampling: (i) ensuring that the active-to-inactive ratio remains approximately equal across both partitions, and (ii) maintaining a training-to-test ratio within a pre-selected range (from 3:1 to 4:1 in this article). This range can be defined by the user via the argument -k: the training-to-test ratio would range from (*k −* 1):1 to *k*:1. For each optimization trial, the maximum pairwise Tanimoto similarity threshold for each class (*τ_A_* for actives and *τ_I_* for inactives) is sampled between 0.00 and a predefined upper limit (*τ*_upper*,A*_ for actives, *τ*_upper*,I*_ for inactives), defaulting to 0.70 for both classes [27, 31, 67, 68, 69, 70], using a Tree-structured Parzen Estimator (TPE) sampler [71]. The predefined upper limits of class-specific thresholds can be easily modified by the user via the arguments -upperActive and -upperInactive. Based on these sampled thresholds, active and inactive instances are clustered separately using a single-linkage clustering method (Supplementary Algorithm S1) [28, 60, 61], ensuring that the maximum Tanimoto similarities across disjoint clusters do not exceed *τ_A_* and *τ_I_*for each respective class. This clustering strategy initially forms preliminary clusters by identifying a list of neighbors for each instance within the given similarity threshold, subsequently linking and merging any clusters that overlap by at least one instance. Following clustering, these groups are allocated into two partitions through *n*_cycle_ (-cycle, defaulting to 1) refinement loops (cf. Allocation strategy). If the resulting partitions satisfy the aforementioned dual constraints, the trial is classified as a SUCCESS and returns the sampled Tanimoto similarity cutoffs as the optimization objectives; otherwise, the trial returns values calculated from a penalty function (Algorithm 1). After a predefined number of optimization trials (*n*_iter_ or -iter), defaulting to 100 as recommended by the Optuna documentation [38], a Pareto front [72] of non-dominated solutions is generated from the successful trials. To select the optimal partition from this Pareto front, our workflow utilizes the DOE score, which quantifies how well actives and inactives are embedded together in a multidimensional physicochemical space [2, 3, 58], as a tie-breaking criterion, favoring the solution that yields the lowest DOE score in the partitioned test set. Consequently, this partitioning mode reduces analog bias, as indicated by the intra-class maximum structural similarity across the two partitions, while also minimizing test-set inter-class bias derived from discrepancies in physicochemical properties between actives and inactives. This establishes a highly challenging scenario, forcing models trained on the training set to generalize to a structurally dissimilar test set containing highly overlapping active and inactive instances within the physicochemical space.

#### Algorithm 1 MolSCISSOR Multivariate Objective Function

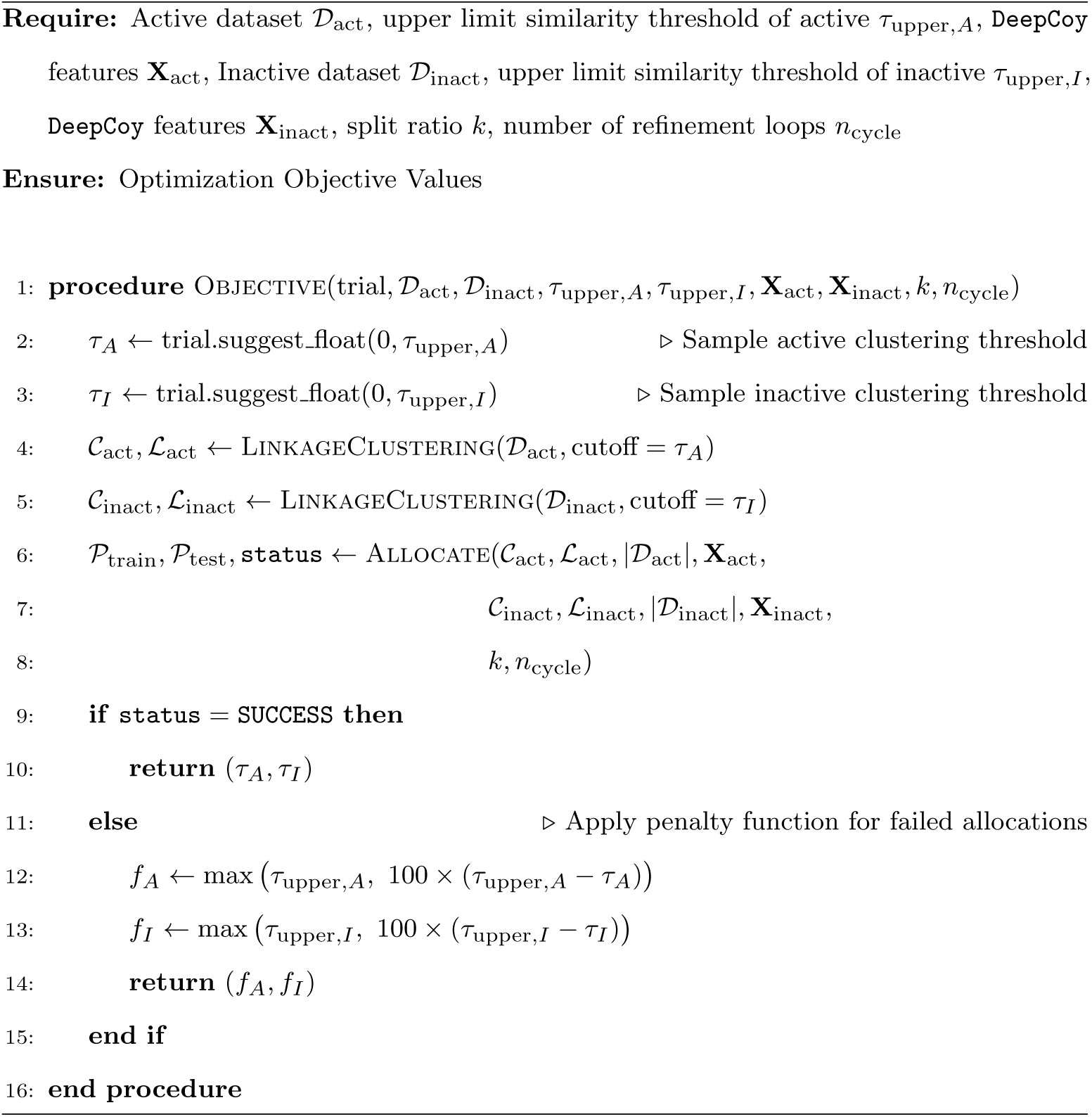

|  |  |  |
|---|---|---|
| **Require:** Active dataset 𝒟<sub>act</sub>, upper limit similarity threshold of active τ<sub>upper,*A*</sub>, DeepCoy features **X**<sub>act</sub>, Inactive dataset 𝒟<sub>inact</sub>, upper limit similarity threshold of inactive τ<sub>upper,*I*</sub>, DeepCoy features **X**<sub>inact</sub>, split ratio *k*, number of refinement loops *n*<sub>cycle</sub> |  |  |
| **Ensure:** Optimization Objective Values |  |  |
| 1: | **procedure** Objective(trial, 𝒟<sub>act</sub>, 𝒟<sub>inact</sub>, τ<sub>upper,*A*</sub>, τ<sub>upper,*I*</sub>, **X**<sub>act</sub>, **X**<sub>inact</sub>, *k*, *n*<sub>cycle</sub>) |  |
| 2: | τ<sub>*A*</sub> ← trial.suggest_float(0, τ<sub>upper,*A*</sub>) | ▷ Sample active clustering threshold |
| 3: | τ<sub>*I*</sub> ← trial.suggest_float(0, τ<sub>upper,*I*</sub>) | ▷ Sample inactive clustering threshold |
| 4: | 𝒞<sub>act</sub>, ℒ<sub>act</sub> ← LinkageClustering(𝒟<sub>act</sub>, cutoff = τ<sub>*A*</sub>) |  |
| 5: | 𝒞<sub>inact</sub>, ℒ<sub>inact</sub> ← LinkageClustering(𝒟<sub>inact</sub>, cutoff = τ<sub>*I*</sub>) |  |
| 6: | 𝒫<sub>train</sub>, 𝒫<sub>test</sub>, status ← Allocate(𝒞<sub>act</sub>, ℒ<sub>act</sub>, \|𝒟<sub>act</sub>\|, **X**<sub>act</sub>, |  |
| 7: | 𝒞<sub>inact</sub>, ℒ<sub>inact</sub>, \|𝒟<sub>inact</sub>\|, **X**<sub>inact</sub>, |  |
| 8: | *k*, *n*<sub>cycle</sub>) |  |
| 9: | **if** status = SUCCESS **then** |  |
| 10: | **return** (τ<sub>*A*</sub>, τ<sub>*I*</sub>) |  |
| 11: | **else** | ▷ Apply penalty function for failed allocations |
| 12: | *f*<sub>*A*</sub> ← max (τ<sub>upper,*A*</sub>, 100 × (τ<sub>upper,*A*</sub> − τ<sub>*A*</sub>)) |  |
| 13: | *f*<sub>*I*</sub> ← max (τ<sub>upper,*I*</sub>, 100 × (τ<sub>upper,*I*</sub> − τ<sub>*I*</sub>)) |  |
| 14: | **return** (*f*<sub>*A*</sub>, *f*<sub>*I*</sub>) |  |
| 15: | **end if** |  |
| 16: | **end procedure** |  |

### 4.3 Allocation strategy

The MolSCISSOR allocation framework, detailed in Algorithm 2, employs an iterative refinement strategy to partition both active and inactive compounds into distinct subsets (*P*_train_ and *P*_test_), taking the disjoint clusters obtained from the aforementioned single-linkage clustering as inputs. The allocation process, comprising normal allocation, 0/1 knapsack-based allocation, and iterative refinement, is guided by stratified sampling-based dual constraints (cf. Core algorithm). A SUCCESS status is returned only when these constraints are satisfied; otherwise, the process terminates with a FAILURE flag.

#### Algorithm 2 MolSCISSOR Allocation

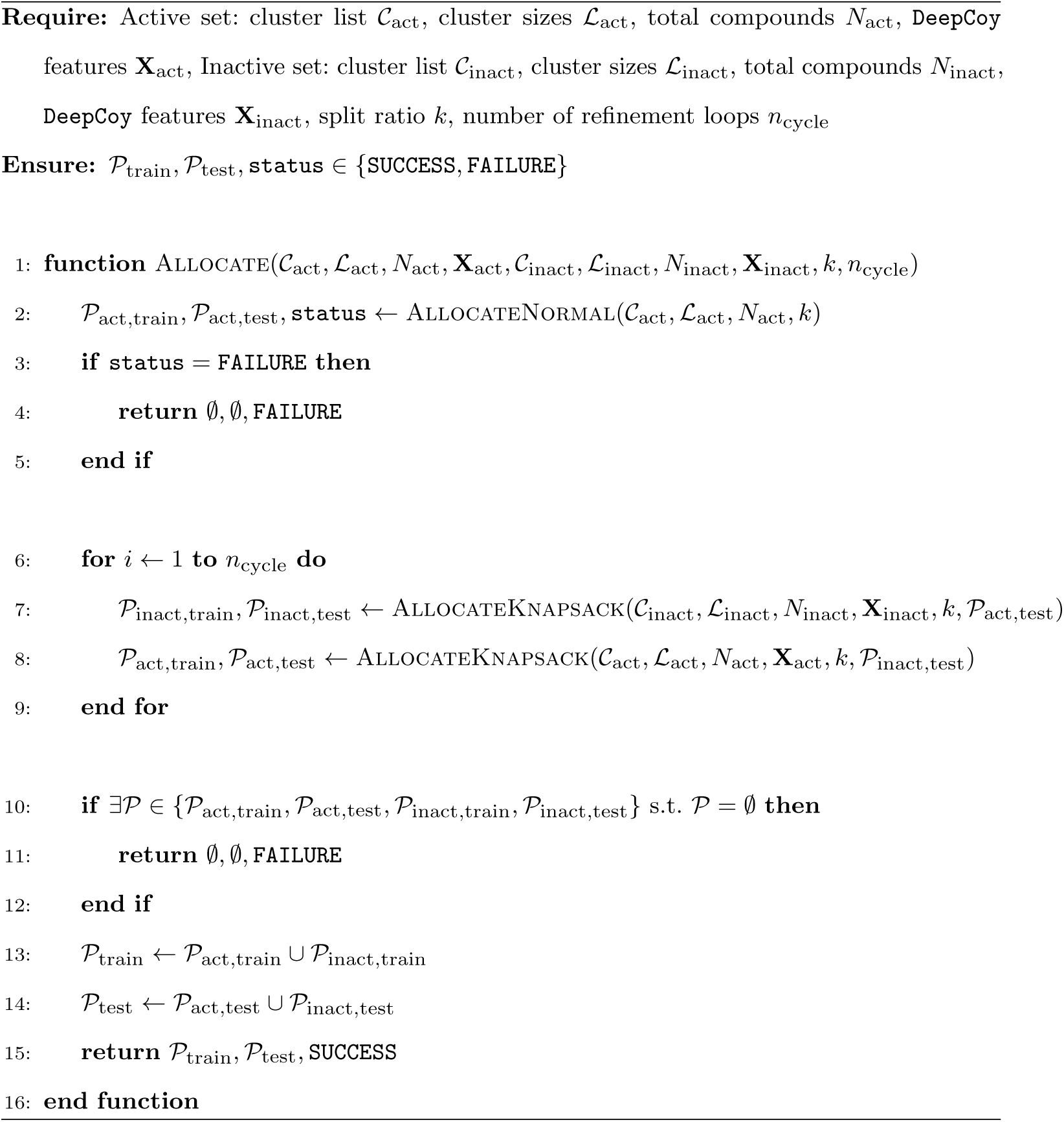

| **Require:** | Active set: cluster list $\mathcal{C}_{\text{act}}$, cluster sizes $\mathcal{L}_{\text{act}}$, total compounds $N_{\text{act}}$, DeepCoy features $\mathbf{X}_{\text{act}}$, Inactive set: cluster list $\mathcal{C}_{\text{inact}}$, cluster sizes $\mathcal{L}_{\text{inact}}$, total compounds $N_{\text{inact}}$, DeepCoy features $\mathbf{X}_{\text{inact}}$, split ratio $k$, number of refinement loops $n_{\text{cycle}}$ |
|---|---|
| **Ensure:** | $\mathcal{P}_{\text{train}}, \mathcal{P}_{\text{test}}, \texttt{status} \in \{\texttt{SUCCESS}, \texttt{FAILURE}\}$ |
| 1: | **function** $\textsc{Allocate}(\mathcal{C}_{\text{act}}, \mathcal{L}_{\text{act}}, N_{\text{act}}, \mathbf{X}_{\text{act}}, \mathcal{C}_{\text{inact}}, \mathcal{L}_{\text{inact}}, N_{\text{inact}}, \mathbf{X}_{\text{inact}}, k, n_{\text{cycle}})$ |
| 2: | $\mathcal{P}_{\text{act,train}}, \mathcal{P}_{\text{act,test}}, \texttt{status} \leftarrow \textsc{AllocateNormal}(\mathcal{C}_{\text{act}}, \mathcal{L}_{\text{act}}, N_{\text{act}}, k)$ |
| 3: | **if** $\texttt{status} = \texttt{FAILURE}$ **then** |
| 4: | **return** $\emptyset, \emptyset, \texttt{FAILURE}$ |
| 5: | **end if** |
| 6: | **for** $i \leftarrow 1$ **to** $n_{\text{cycle}}$ **do** |
| 7: | $\mathcal{P}_{\text{inact,train}}, \mathcal{P}_{\text{inact,test}} \leftarrow \textsc{AllocateKnapsack}(\mathcal{C}_{\text{inact}}, \mathcal{L}_{\text{inact}}, N_{\text{inact}}, \mathbf{X}_{\text{inact}}, k, \mathcal{P}_{\text{act,test}})$ |
| 8: | $\mathcal{P}_{\text{act,train}}, \mathcal{P}_{\text{act,test}} \leftarrow \textsc{AllocateKnapsack}(\mathcal{C}_{\text{act}}, \mathcal{L}_{\text{act}}, N_{\text{act}}, \mathbf{X}_{\text{act}}, k, \mathcal{P}_{\text{inact,test}})$ |
| 9: | **end for** |
| 10: | **if** $\exists \mathcal{P} \in \{\mathcal{P}_{\text{act,train}}, \mathcal{P}_{\text{act,test}}, \mathcal{P}_{\text{inact,train}}, \mathcal{P}_{\text{inact,test}}\}$ s.t. $\mathcal{P} = \emptyset$ **then** |
| 11: | **return** $\emptyset, \emptyset, \texttt{FAILURE}$ |
| 12: | **end if** |
| 13: | $\mathcal{P}_{\text{train}} \leftarrow \mathcal{P}_{\text{act,train}} \cup \mathcal{P}_{\text{inact,train}}$ |
| 14: | $\mathcal{P}_{\text{test}} \leftarrow \mathcal{P}_{\text{act,test}} \cup \mathcal{P}_{\text{inact,test}}$ |
| 15: | **return** $\mathcal{P}_{\text{train}}, \mathcal{P}_{\text{test}}, \texttt{SUCCESS}$ |
| 16: | **end function** |

### 4.3.1 Normal allocation

The workflow begins by generating an initial test partition for the active molecules through the AllocateNormal sub-process (Supplementary Algorithm S2), which filters and groups compounds according to their pre-calculated cluster sizes. The process starts by setting upper and lower cluster size boundaries. In detail, to prevent single massive clusters from unbalancing the data split, the algorithm evaluates large-(*L*[*i*] *> lim*_upper_) and medium-sized (*lim*_lower_ *< L*[*i*] *≤ lim*_upper_) clusters first; if any cluster exceeds the upper boundary, or if their cumulative size surpasses partition capacity limits, the process immediately terminates and returns a FAILURE flag. For the remaining small clusters (*L*[*i*] *≤ lim*_lower_), the algorithm sorts them in descending order and accumulates them into *P*_test_ until the total count reaches the lower boundary threshold. The remaining unassigned compounds are then distributed to *P*_train_ as the complement set. This sub-process concludes with a final verification check to ensure the size of the resulting partitions complies with the stratified sampling-based dual constraints.

### 4.3.2 0/1 knapsack-based allocation and iterative refinement

Following the initialization with a SUCCESS status via AllocateNormal, the framework enters a refinement loop for a predefined number of cycles (*n*_cycle_) using the knapsack-based allocation AllocateKnapsack (Supplementary Algorithm S3). This sub-process handles the iterative refinement phase by framing subset selection as a classic 0/1 knapsack problem introduced by Martello and Toth [59]. The algorithm first establishes a weight capacity threshold derived from the total compound count *N* and the split ratio *k*, which controls the training-to-test ratio, then immediately skips any cluster that exceeds this limit. For the remaining valid clusters, the framework computes an optimization profit based on the spatial distance between their 26-dimensional DeepCoy physicochemical descriptor vectors and the fixed reference set (*D*_ref_). A dynamic programming solver (KnapsackSolverDP) is then deployed to determine the optimal configuration of clusters that maximizes profit without surpassing the capacity threshold. This solver is implemented from the Google OR-Tools library (v.9.15.6755) using the KNAPSACK DYNAMIC PROGRAMMING SOLVER backend initialized as “ClusterSelection”.

Within each iteration, this knapsack-based allocation reciprocally alternates between the active and inactive compound spaces. Specifically, the algorithm dynamically updates the reference set (*D*_ref_) by using the newly optimized subset of one compound pool (e.g., the active subset) to guide the re-allocation of the other (e.g., the inactive subset). This back-and-forth optimization scheme runs for *n*_cycle_ steps to allow both subsets to progressively update. This mechanism systematically minimizes the spatial embedding mismatch between the final test subsets, effectively mitigating inter-class bias caused by physicochemical property disparities across the chemical feature space in the test set. Ultimately, the final selected clusters are allocated into *P*_test_, while the remaining unselected compounds constitute *P*_train_ via complement set subtraction.

### 4.4 bblean clustering-based split

The clustering process utilizing the bblean algorithm followed the same protocol as described in its GitHub repository (https://github.com/mqcomplab/bblean, accessed September 2026) [36, 37]. First, input files comprising canonicalized and validated SMILES strings for active and inactive sets were used to generate packed Morgan fingerprints (2,048 bits, radius 2) via the bb fps-from-smiles command, and the outputs were saved in .npy format. Next, from these outputs, the optimal similarity threshold controlling the cluster radius was estimated using the bb thresh command. The bblean clustering was then executed with this optimal threshold value while keeping all other parameters at their defaults, yielding disjoint clusters. To ensure a fair comparison with our approach, these clusters were assigned into either training or test partition with the exact same allocation strategy as that of MolSCISSOR, comprising normal allocation, 0/1 knapsack-based allocation, and iterative refinement (cf. Allocation strategy).

### 4.5 Baseline model benchmarking

To evaluate the difficulty of the VS task posed by each LIT-PCBA variant (AVE-unbiased LIT-PCBA, bblean-LIT-PCBA, and SPLIT-PCBA), we trained target-specific ML models on each training partition and evaluated their predictive performance on the corresponding test set. For this evaluation, we employed three classical ML algorithms (RF, XGB, and SVC), a deep neural network baseline (MLP), a graph neural network (GraphConv), and a fine-tuned pre-trained transformer-based foundation model (ChemBERTa), each with three different seeds to ensure reproducibility and robustness of the evaluation framework.

The ML-based models were implemented using the RandomForestClassifier and SVC classes from the sklearn.ensemble and sklearn.svm modules (scikit-learn v.1.7.2) for RF and SVC, respectively, and the XGBClassifier class from the xgboost library (v.3.2.0) for XGB. All hyperparameters were kept at their default values. For MLP, we utilized the architecture implemented in DeepPurpose [39], a deep learning library for drug properties prediction. The network consists of two main components: (i) a feature encoder that maps the input vector through hidden layers of 1024, 256, and 64 units to yield a 256-dimensional latent representation; and (ii) a downstream classifier head comprising three linear layers with 1024, 1024, and 512 units, using ReLU activations and dropout (*p* = 0.1), followed by a sigmoid output layer. For these models, two types of input features were employed. First, we considered three classes of molecular fingerprints (each with 2,048 bits): circular Morgan fingerprints (radius 2, generated via the GetMorganFingerprintAsBitVect function from the rdkit.Chem.AllChem module), path-based Daylight fingerprints (via the FingerprintMols.FingerprintMol function from the rdkit.Chem.Fingerprints module), and torsion-based fingerprints (via the GetHashedTopologicalTorsionFingerprintAsBitVect function from the rdkit.Chem.rdMolDescriptors module). Second, 217 RDKit physicochemical descriptors were utilized, which were standardized using a StandardScaler fitted on the training set and subsequently applied to the test set.

For the GraphConv model, compounds were first featurized from SMILES strings using the ConvMolFeaturizer class implemented in DeepChem (v.2.5.0) [40]. Each heavy atom was encoded as a 75-dimensional node feature vector (detailed in Supplementary Table S3), while covalent bonds were stored as canonical undirected adjacency lists. The architecture was constructed using GraphConvModel from DeepChem, comprising two sequential graph convolutional message-passing layers, followed by a dense layer of 2,048 dimensions and a final classification output head. All graph models were executed in PyTorch (v.2.6.0+cu124) with CUDA 12.4 acceleration.

For ChemBERTa, we utilized the ChemBERTa-77M-MTR transformer backbone (https://huggingface.co/DeepChem/ChemBERTa-77M-MTR, accesed October 2026) [41], a chemical foundation model based on the RoBERTa architecture pre-trained on 77 million unique PubChem SMILES strings using a Multi-Task Regression (MTR) objective over 200 calculated physicochemical properties. Canonical SMILES strings were tokenized and truncated to a maximum sequence length of 512 tokens to match the pre-training positional embedding capacity. The model was fine-tuned on the canonicalized training set using sequence classification head initialization with default Hugging Face fine-tuning hyperparameters (learning rate of 5 *×* 10*^−^*^5^, three epochs), after which inference was evaluated on the test set.

### 4.6 Performance evaluation

The screening performance of the trained target-specific ML models, comprising baseline RF, XGB, SVC, MLP, GraphConv, and ChemBERTa architectures, evaluated on the corresponding test sets was quantified using three metrics: ROC-AUC, NEF1%, and BEDROC(*α* = 160.9) [27, 28, 42, 73].

To complement the evaluation of the training-test splitting frameworks, the underlying biases of the resulting training-test pairs of each LIT-PCBA variant were assessed using the following metrics (all similarity calculations were performed using Tanimoto similarity based on Morgan fingerprints, 2,048 bits, radius 2):

i. the maximum intra-class Tanimoto similarity between the training and test partitions:

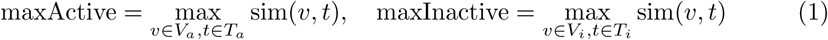

where *T_a_*, *T_i_*, *V_a_*, *V_i_* are training actives, training inactives, test actives, and test inactives, respectively;
ii. (ii) the AVE bias [11]:

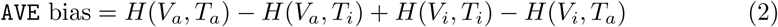

where 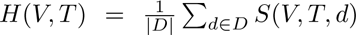 denotes the cumulative nearest neighbor function across discrete distance thresholds *d* ɛ *D =* [0, 1], with 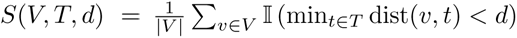; and
iii. (iii) the scaled DataSAIL information “leakage” L(*π*) [35]:

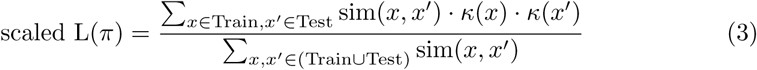

where *κ*(*x*) is the cardinality of data points or clusters. In this case, *κ*(*x*) = *κ*(*x^′^*) = 1 since elementary data points are considered.

A lower maximum intra-class Tanimoto similarity indicates greater structural dissimilarity between same-class compounds across the two partitions, preventing model overfitting via structural memorization. Similarly, a low AVE bias minimizes trivial structural clumping [11], and a low scaled L(*π*) indicates reduced cross-split information leakage and sampling bias [35], thereby ensuring a more rigorous and unbiased evaluation of model generalization.

### 4.7 Statistical analysis

Statistical analysis was conducted to ensure rigorous and reliable comparisons across methods. Two-sided Mann-Whitney U tests with Bonferroni correction were performed to compare MolSCISSOR against the other two partitioning procedures. Besides, one-sample two-sided Wilcoxon signed-rank tests were performed to compare the ROC-AUC values of the evaluated models with the random-guessing baseline of 0.5. All statistical tests were executed using the scipy package (v.1.17.1).

### 4.8 Benchmarking memory footprints

To evaluate practical utility, the memory efficiency of five partitioning tools was compared: the AVE method, the bblean clustering-based split, MolSCISSOR, and two recently published advanced tools (Lo-Hi [34] and DataSAIL [35]). To ensure a fair comparison, the peak memory usage of all tools was recorded under identical execution conditions: a single CPU core, one worker process, and a memory cap of 200 GB RAM per task. Default execution parameters were maintained across all runs: (i) for the AVE method, a maximum of 300 optimization iterations (-maxIter 300) with the convergence criterion set to an AVE bias below 0.01; (ii) for bblean, default parameters including a branching factor of 254 and a diameter merging criterion; (iii) for MolSCISSOR, one refinement loop (-cycle 1) and 100 Optuna optimization iterations (-iter 100); (iv) for the Lo-Hi splitter, default similarity threshold of 0.4, train min frac of 0.75, and test min frac of 0.2; and (v) for DataSAIL, a training-to-test ratio of 0.8:0.2 with C1e technique using single-linkage clustering based on ECFP4 fingerprints. Computational memory efficiency was quantified by recording the peak memory consumption of each task via the SLURM job efficiency utility (seff). Target datasets were categorized by size into small (*<*10,000 compounds), medium (10, 000 to 100, 000 compounds), and large (*>*100,000 compounds).

## 5 Data availability

The AVE-unbiased LIT-PCBA benchmark was downloaded from https://drugdesign.unistra.fr/LIT-PCBA in March 2026. The raw data and results reported in this study are available through our GitHub repository (https://github.com/caominhtr/molscissor) and Zenodo (https://doi.org/10.5281/zenodo.23942580).

## 6 Code availability

The scripts required to run BitBIRCH-Lean, Lo-Hi, and DataSAIL were obtained from their GitHub repositories (https://github.com/mqcomplab/bblean, https://github.com/steshinss/lohi_splitter, and https://github.com/kalininalab/DataSAIL, respectively) and used without modification. The source code for MolSCISSOR is available through our GitHub repository (https://github.com/caominhtr/molscissor).

## 8 Authors’ contributions

V.-K.T.-N. conceived the research, defined the study design, and supervised the project, with contributions from I.V.T., P.J.B., and O.T. C.-M.T. was responsible for developing the MolSCISSOR code, conducting dataset partitioning, performing the subsequent evaluation, analysis and interpretation of the results under the supervision of V.-K.T.-N. V.-K.T.-N. and C.-M.T. drafted the initial manuscript, with input from I.V.T., P.J.B., and O.T. All authors reviewed, revised, and approved the final version of the manuscript.

